# Female fruitflies use gustatory cues to exhibit reproductive plasticity in response to the social environment

**DOI:** 10.1101/2021.03.10.434778

**Authors:** E.K. Fowler, S. Leigh, W.G. Rostant, A. Thomas, A. Bretman, T. Chapman

## Abstract

Animals can exhibit remarkable reproductive plasticity in response to their social surroundings, with profound fitness consequences. The study of such plasticity in females, particularly in same-sex interactions, has been severely neglected. Here we measured the impact of variation in the pre-mating social environment on reproductive success in females and tested the underlying mechanisms involved. We used the *Drosophila melanogaster* model system to test the effect of varying female group size prior to mating and deployed physical and genetic methods to manipulate the perception of different social cues and sensory pathways. We found that socially isolated females were significantly more likely to retain unfertilised eggs before mating, but to show the opposite pattern and lay significantly more fertilised eggs in the 24h after mating, in comparison to grouped females. More than 48h of exposure to other females was necessary for this socially-induced plasticity to be expressed. Neither olfactory nor visual cues were involved in mediating these responses. Instead, we found that females detected other females through direct contact with the deposits they leave behind, even in the absence of eggs. The results demonstrate that females show striking reproductive plasticity in response to their social surroundings and that the nature of their plastic reproductive responses, and the cues they use, differ markedly from those of males. The results emphasise the stark contrasts in how each sex realises reproductive success.

## Introduction

Phenotypic plasticity (the expression of different phenotypes from the same genotype) is a widespread and important component of fitness, allowing individuals to adaptively alter their behaviour or physiology in response to environmental variation (Pigliucci, 2001; West-Eberhard, 2003). An organism’s social surroundings (e.g. the local density and ratio of male and female conspecifics and heterospecifics) can vary considerably (Kasumovic & Brooks, 2011). Sex differences in birth and death rates or sexual maturity can cause temporal shifts in sex ratio, either on an immediate, short-term basis or over seasons or successive years. Other factors such as immigration, dispersal and the level of predation also contribute to a dynamic social environment (Kasumovic & Brooks, 2011). The density and identity of individuals in the social milieu can signal resource quality or the expected likelihood of competition (Davis *et al*., 2011). For example, the sex ratio of conspecifics could indicate the level of competition for mating opportunities, or for sex-specific resources such as oviposition sites. Detection of information from heterospecifics may also be beneficial if habitat requirements overlap between species. If this is the case, the overall density of individuals, independent of species, could signal expected levels of nutrient availability or quality, predation risk (Huang *et al*., 2011) or oviposition sites. Given that variation in the social environment has significant consequences for the level of reproductive competition or resource availability, individuals with the ability to detect cues from their social environment and adjust their phenotype accordingly can increase their fitness (Bretman *et al*., 2013).

The effect of the social environment on phenotypic plasticity in males has been well studied in the context of sperm competition (Bretman *et al*., 2011; Dore *et al*., 2018; Parker & Pizzari, 2010; Wedell *et al*., 2002). *Drosophila melanogaster* fruitflies in particular have proved to be a valuable model in this context. Males can precisely and flexibly adjust their ejaculate composition and extend copulation duration in response to the presence of conspecific rival males (Bretman *et al*., 2011; Bretman *et al*., 2013; Garbaczewska *et al*., 2013; Wigby *et al*., 2009). These plastic adjustments enable males to secure a greater share of the paternity when sperm competition is perceived to be high, while conserving costly resources when sperm competition is unlikely (Bretman *et al*., 2009).

Despite extensive studies into male social plasticity, we know very little about the corresponding context in females – i.e. whether and how they might adjust their reproductive output in response to the intrasexual environment. Naïve females can exhibit social learning and adjust their oviposition site preferences to match those of experienced mated females (Sarin & Dukas, 2009) and oviposition preference can be influenced both by pheromonal cues from conspecifics (Dumenil *et al*., 2016; Malek & Long, 2020; Wertheim *et al*., 2002) and the presence of predators (Kacsoh *et al*., 2015). Female social plasticity has also been considered in the context of mate choice and differential responses to male characteristics (Bailey & Zuk, 2008; Billeter *et al*., 2012; Filice & Long, 2017; Fox *et al*., 2019). However, whether females can plastically optimise their reproductive output according to the general expectation of reproductive or resource competition (e.g. as signalled by the presence of other females) is not yet known and remains an important and unanswered question.

For fitness benefits of phenotypic plasticity to be accrued by either sex, and plasticity itself to evolve, mechanisms for the accurate perception of cues that reliably indicate the social or sexual environment are required. In male *D. melanogaster* cues of competition are detected via multiple, interchangeable olfactory, auditory and tactile sensory pathways (Bretman *et al*., 2011). This multimodal strategy is predicted to decrease the risk of costly mismatches between environment and phenotype in highly variable environments (Dore *et al*., 2018) enabling males to accurately perceive information on the species, sex and prevalence of other individuals, and respond appropriately to the level of sperm competition (Bretman *et al*., 2017). Whether females deploy any such multimodality via complex cues is also not yet known.

Here, we address these omissions by testing the hypothesis that *D. melanogaster* females plastically adjust their reproductive investment according to the con and hetero - specific intrasexual social environment. Focal females were either housed in isolation or with three other females before being given the opportunity to mate with a single male. We recorded mating times and the number of eggs (fecundity) laid in the 3 days before and in the 24h after mating. During the social exposure phase, all females were virgins. This allowed us to test the response of females to the same sex environment without the confounding effects of previous mates or male pheromones. We thus investigated the effect of the proximate social environment on both virgin egg laying, and subsequent post-mating fecundity. We also probed the underpinning mechanisms involved by varying social exposure time and by restricting the perception of social cues by using genetic and physical manipulations.

## Results

### Female fecundity responses to variation in the social environment and effect of exposure to con- vs hetero-specific females

We measured the impact of pre-mating social isolation versus exposure to other females on the reproductive output of focal *D. melanogaster* females following a single mating. Virgin focal females were exposed to different social environments for 72h prior to mating, and fecundity was measured as the number of eggs laid in the 24h period following mating. During the post-mating period, focal females previously held in groups of four conspecifics laid significantly fewer eggs than previously socially isolated females (Figure 1a, *F*_(1, 84)_ = 4.48, *p* = 0.037). Similarly, *D. melanogaster* females held with three heterospecific females (either *D. simulans* or *D. yakuba*) prior to mating were also significantly less fecund following mating than were socially isolated females (*simulans*: *F*_(1, 76)_ = 4.64, p = 0.035; *yakuba*: *F*_(1, 90)_ = 18.00, *p* = 5.36 × 10^−5^) (Figure 1b).

**Figure 1.**
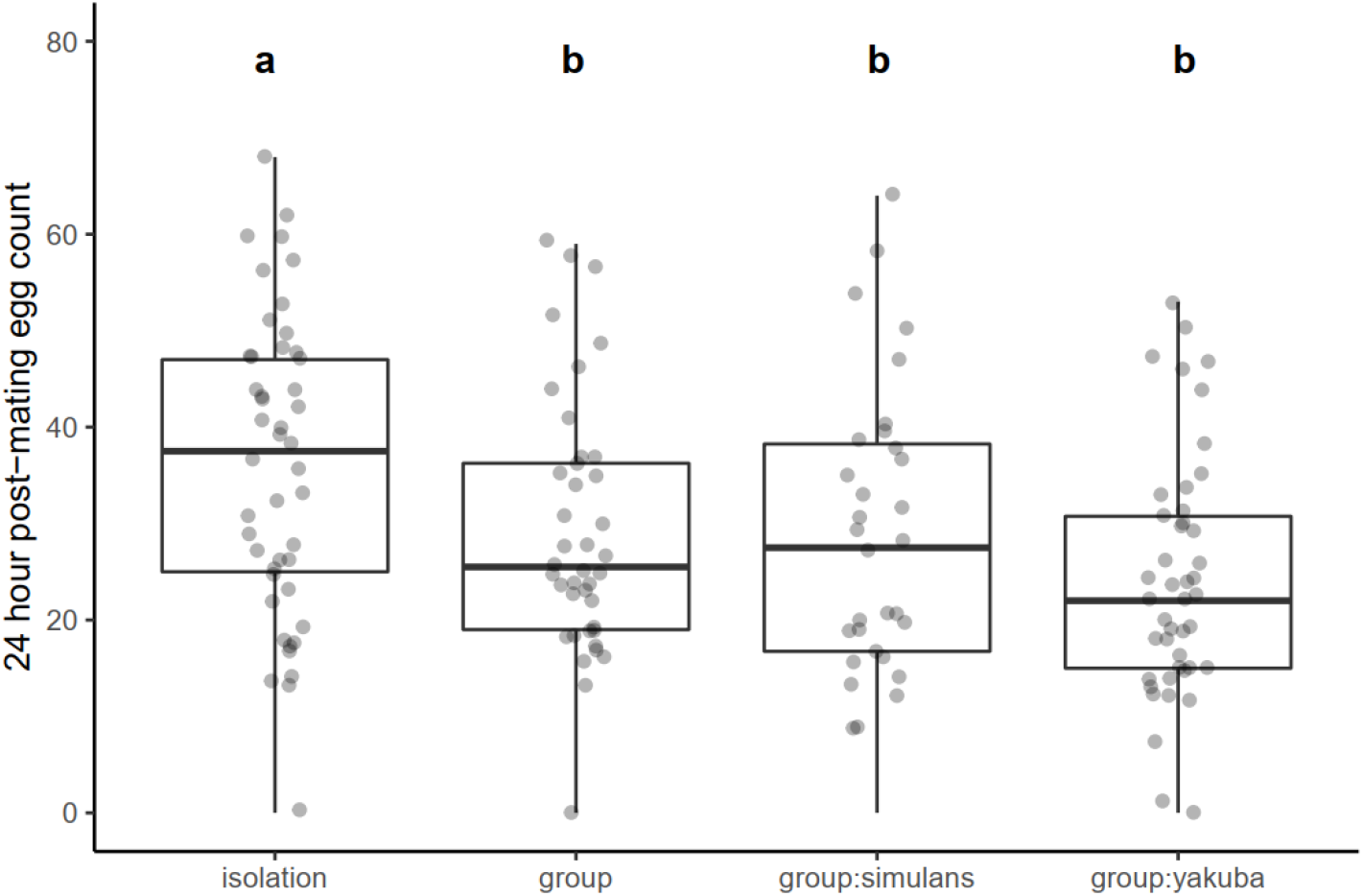
*D. melanogaster* females exposed to con- or hetero-specific females prior to mating show significantly decreased post-mating fecundity. *D. melanogaster* females were kept socially isolated (‘isolation’) or exposed to con- (‘group’) or hetero-specific females (‘group:simulans’ or ‘group:yakuba’) for 72h prior to mating. Fecundity was measured as the number of eggs laid by each female in the 24h period following mating. Boxplots show interquartile range (IQR) and median in the box, and whiskers represent the largest and smallest values within 1.5 times the IQR above and below the 75 ^th^ and 25^th^ percentiles, respectively. Raw data points are plotted with jitter. Treatments not sharing a letter are significantly different from one another (p < 0.05).

### Effect of length of social exposure period on post-mating fecundity

The response of *D. melanogaster* female fecundity to the pre-mating social environment was affected by the length of exposure to conspecific females. When focal females were exposed to the different social environment treatments for 2, 4, 8, 24, 48 or 72h prior to mating, only those exposed for 72h showed a significant reduction in fecundity compared to isolated females (*F*_(1, 120)_ = 20.85, *p* = 1.21 × 10^−5^). The effect of social treatment on eggs was marginally non-significant for the 48h exposure period (*F*_(1, 115)_ = 3.68, *p* = 0.058), and not significant for all other shorter periods (2h: *F*_(1, 87)_ = 0.80, *p* = 0.37; 4h: *F*_(1, 86)_ = 0.03, *p* = 0.87; 8h: *F*_(1, 75)_ = 1.28, *p* = 0.26; 24h: *F*_(1, 115)_ = 0.30, *p* = 0.59) (Figure 2).

**Figure 2.**
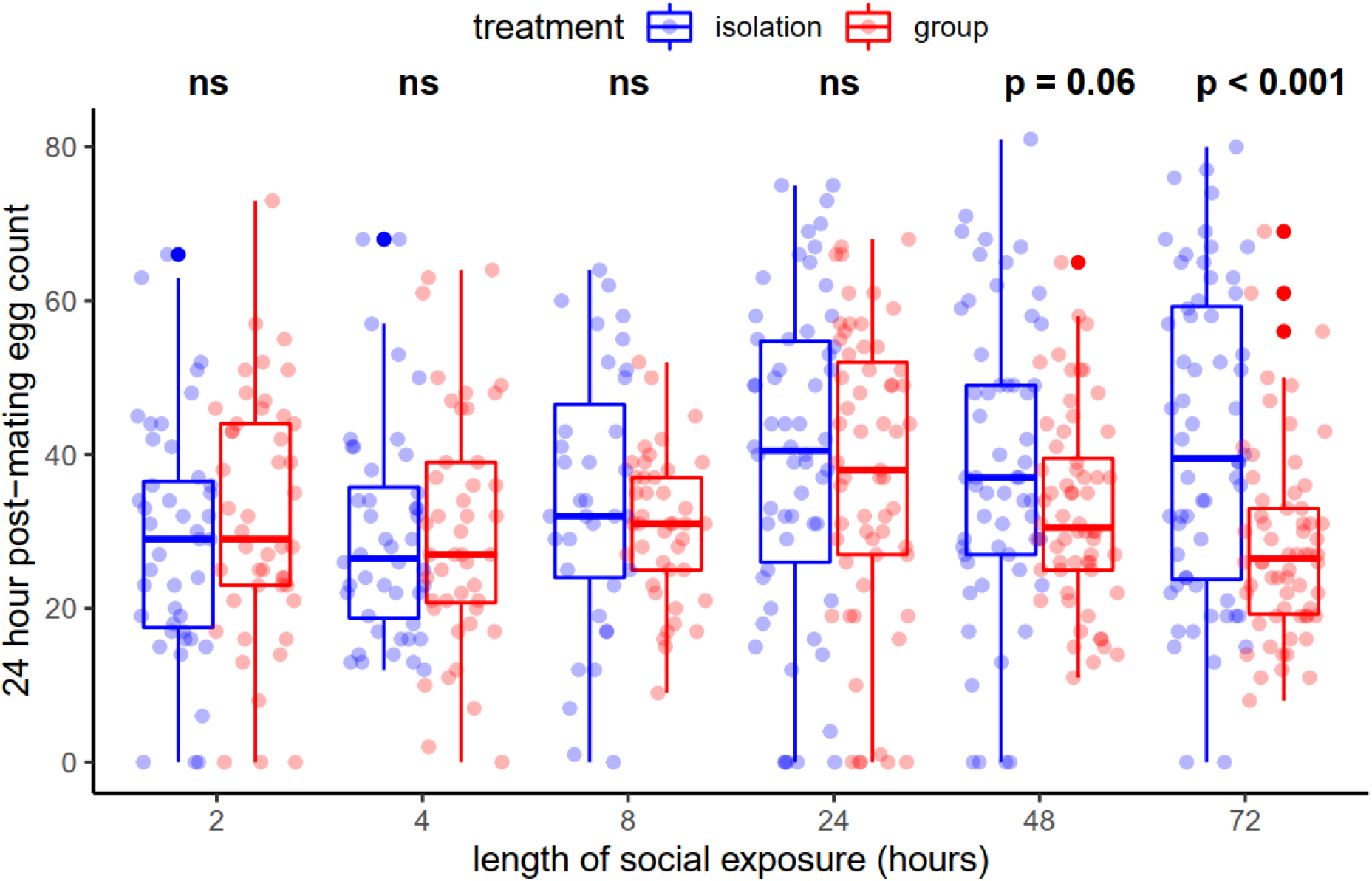
*D. melanogaster* females require 72h of exposure to conspecifics to express fecundity plasticity. Females were housed in ‘isolation’ (blue) or in ‘group’ (red boxes) treatments, for between 2h and 72h prior to mating. Fecundity was measured as the number of eggs laid in the 24h period following mating. Statistical significance indicated above box pairs (ns: p < 0.1). Boxplots as in Figure 1.

### Investigation of whether exposure to eggs or to female deposits in the absence of eggs are required for social exposure effects on post-mating fecundity

To identify the cues that *D. melanogaster* females use to respond to the presence of others, we analysed whether a female’s post-mating fecundity responded to the physical presence of other females, to their eggs or to the deposits they leave behind even in the absence of egg laying. We compared the post-mating fecundity of females subjected to the following treatments: ‘isolation’, ‘group’, ‘group - eggless females’, ‘isolation - female deposits’, ‘isolation - egg-spiked’. Consistent with the previous experiments, ‘group’ females laid significantly fewer eggs than females from the ‘isolation’ treatment (*OvoD1* control: *F*_(1, 81)_ = 26.40, *p* = 1.88 × 10^−6^ (Figure 3A); egg-spiked control: *F*_(1, 76)_ = 20.45, *p* = 2.22 × 10^−5^ (Figure 3B)). Furthermore, females from the ‘group - eggless females’, ‘isolation - female deposits’, and ‘isolation - egg-spiked’ treatments also laid significantly fewer eggs in comparison to females from the ‘isolation’ treatment (deposits: *F*_(1, 88)_ = 8.20, *p* = 0.0052; eggless: *F*_(1, 77)_ = 4.29, *p* = 0.042 (Figure 3A); egg-spiked: *F*_(1, 69)_ = 7.11, *p* = 0.0010 (Figure 3B)).

**Figure 3.**
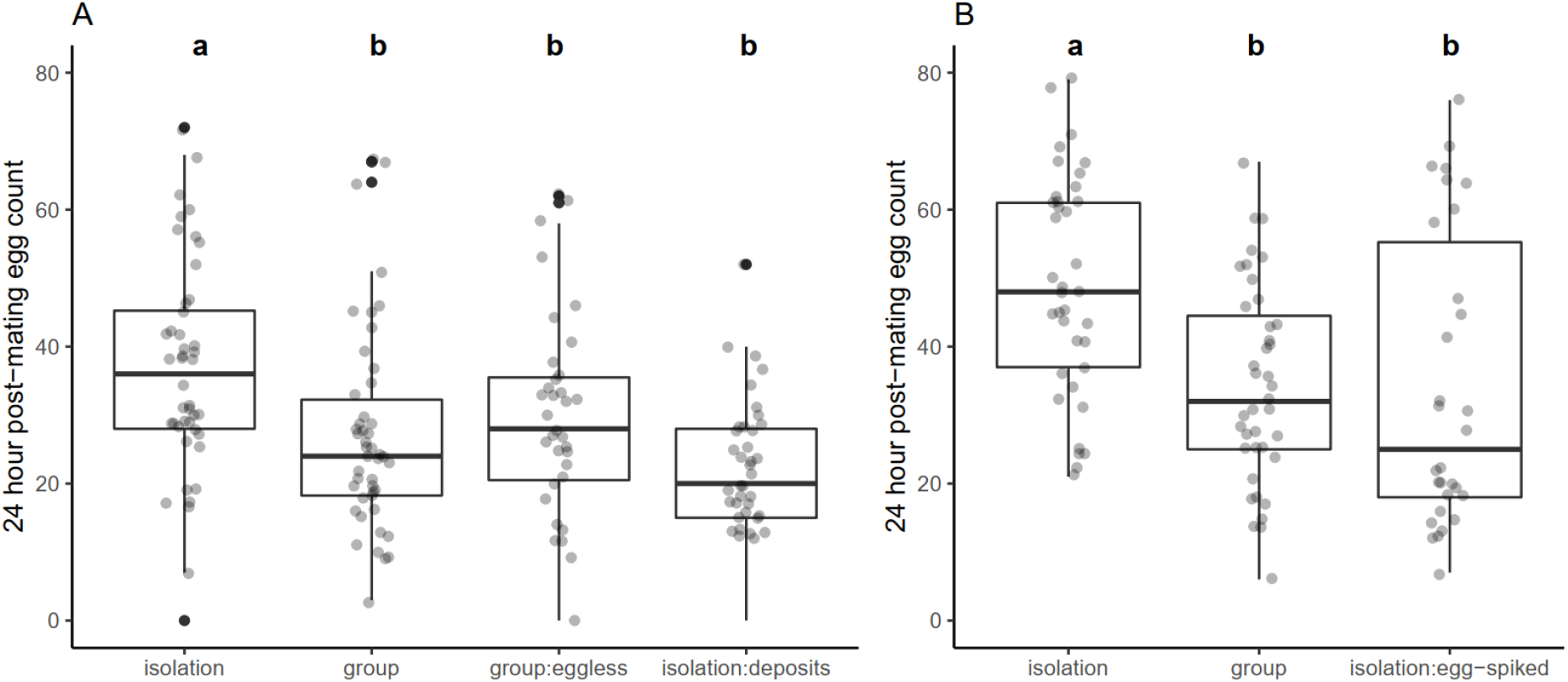
*D. melanogaster* females respond to their social environment by detecting the deposits left by other females, even in the absence of eggs. (A) Wildtype focal females were either isolated in clean vials (‘isolation’), housed in groups of four in clean vials (‘group’), housed with three *OvoD1* females (‘group:eggless’) or housed in vials previously occupied by three *OvoD1* females (‘isolation:deposits’). (B) Wildtype focal females housed in isolation, in groups of four or in vials containing eggs laid by previous wildtype occupants (‘isolation:egg-spiked’). Fecundity was measured as the number of eggs laid by the focal female in the 24h period following a single mating. Boxplots as in Figure 1. Within each plot, treatments not sharing a letter are significantly different from one another (p < 0.05).

### Investigation of the sensory pathways required to detect cues of social exposure effects on post-mating fecundity

To identify the sensory pathways used by focal females to detect the cues contained within female deposits identified as important above, we restricted olfactory, tactile/gustatory and visual inputs. Each sensory input test included socially isolated and group control treatments. In the olfactory restriction experiments, antennaless females laid significantly fewer eggs in the group versus isolation treatment (*F*_(1, 62)_ = 6.43, *p* = 0.014), consistent with the unmanipulated controls (though in this control the group versus isolation comparison was marginally non-significant (*F*_(1, 83)_ = 3.58, *p* = 0.062; Figure 4a). Antennal removal only partially restricts olfactory sensory pathways, since a secondary olfactory system is located in the maxillary palps which thus remained intact (Laissue & Vosshall, 2008). Therefore, to restrict olfactory senses more precisely, we complemented the antennal removal experiment by testing the responses of focal females with a knockout mutation in the broadly expressed olfactory receptor, Orco, which is associated with volatile pheromone sensing (Larsson *et al*., 2004). As with antennaless females, *Orco* knockout females maintained significant fecundity responses to their social environment comparable with those of wild type controls (*Orco*: *F*_(1, 66)_ = 5.13, *p* = 0.027, control: *F*_(1, 88)_ = 4.22, *p* = 0.043; Figure 4b).

**Figure 4.**
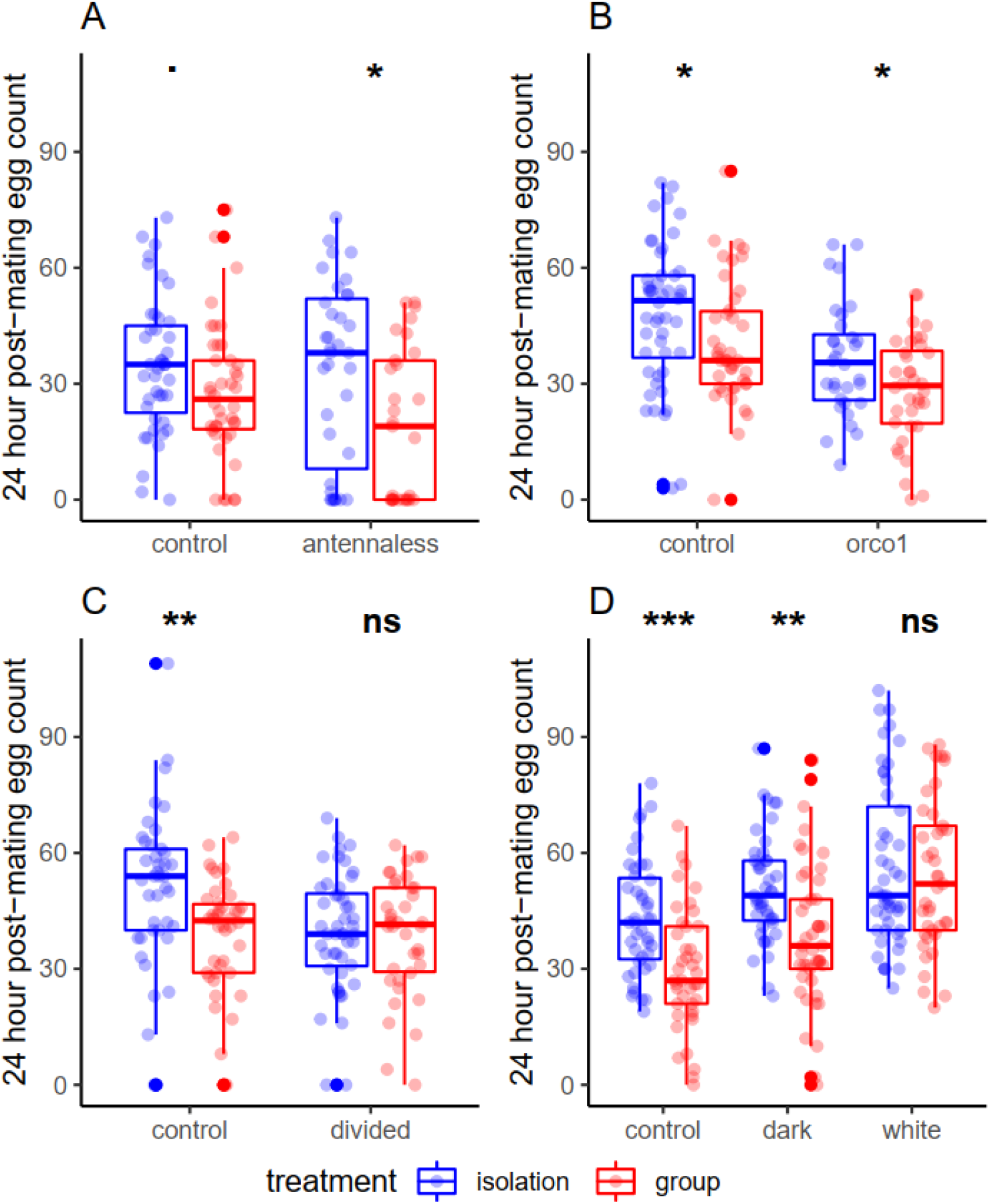
*D. melanogaster* females respond to their social environment by using tactile / gustatory sensory pathways. (A) Olfactory restriction through antennal removal. Intact focal females (‘control’) and olfactory-manipulated focal females with no third antennal segment (‘antennaless’) were kept in isolation or in a group with three intact non-focal females. (B) Olfactory restriction through *Orco* knockout. Wildtype Dahomey females (‘control’) or females lacking the general olfactory receptor Orco (‘*orco*^*1*^’) were kept in isolation or in a group with three Dahomey non-focal females. (C) Tactile/gustatory restriction. Focal females were housed in a standard vial (‘control’) or in a vial with a transparent, perforated divide (‘divided’). For the divided group treatment, focal females were physically separated from the three non-focals by the divide. (D) Visual restriction. Wildtype females held under standard light conditions (‘control’), wildtype females held in darkness (‘dark’) and *white* females (‘white’) were kept in isolation or exposed to three wildtype non-focal females. Fecundity was measured as the number of eggs laid in the 24h period following mating. Boxplots as in Figure 1.

In tests of tactile and gustatory cues, focal females were separated from non-focals in the same vial using a perforated acetate divide. When direct contact with other females was restricted in this way, there was no significant difference in fecundity between grouped and isolated females (*F*_(1, 84)_ = 0.05, *p* = 0.82), in contrast to the control (*F*_(1, 81)_ = 9.31, *p* = 0.0031; Figure 4c).

To manipulate visual input cues, we used either wild-type focal females held in darkness throughout the social exposure period, or vision-defective *white* focal females held under normal conditions (Ferreiro *et al*., 2018). Females held in darkness showed the same significant fecundity responses to social environment as did the control (darkness: *F*_(1, 86)_ = 11.56, *p* = 0.001; control: *F*_(1, 82)_ = 15.97, *p* = 1.40 × 10^−4^; Figure 4d). In contrast, *white* focal female fecundity was unaffected by social environment (*white*: *F*_(1, 87)_ = 0.21, *p* = 0.65; Figure 4d).

### Effect of social environment on virgin egg retention

To test for any potential associations of pre- and post-mating fecundity plasticity we also examined the number of eggs laid by isolated and grouped females prior to mating. Eggs laid by the focal female in the group treatment were distinguished from those of the non-focal by dyeing non-focal females with Sudan Red. Thus focal eggs were white and non-focal eggs were pink. We analysed the egg count data in two steps. First, we split the data into two groups – ‘layers’ (≥ 1 egg laid by focal) or ‘retainers’ (zero eggs laid by focal) and compared the likelihood of focal females from the two social treatments to lay at least one egg. Second, we excluded all zero-counts from the data and compared the numbers of eggs laid by ‘layers’ between the social treatments. For days 1 and 3 of social exposure, isolated females were significantly more likely to retain virgin eggs (i.e. lay zero eggs) than were grouped females (day 1: *X*^2^_1_ = 17.8, *p* = 2.43e-05; day 3: *X*^2^_1_ = 11.5, *p* = 0.0007; Table S2). There was no significant difference on day 2 (*X*^2^_1_= 1.3, *p* = 0.26). Combining data across the 72h period, isolated females were more likely to retain their eggs than were grouped females (*X*^2^_1_= 12.2, *p* = 0.00048; Figure 5a). Of the ‘layers’, isolated females laid significantly more eggs on day 1 than did grouped females (*F*_(1, 53)_= 6.31, *p* = 0.015). However, egg counts did not vary significantly with social treatment on days 2 or 3 or when all days were combined (day 2: *F*_(1, 35)_ = 1.98, *p* = 0.17; day 3: *F*_(1, 40)_ = 0.74, *p* = 0.39; combined: *F*_(1, 67)_ = 0.13, *p* = 0.72; Figure 5b). Analysis of the fecundity of these same females after mating showed that, consistent with previous experiments, grouped females laid significantly fewer eggs post - mating than did isolated females (*F*_(1, 86)_ = 13.35, *p* = 4.43 × 10^−4^; Figure S1). In both social treatments, there was a negative relationship between the number of pre - and post-mating eggs laid (isolation: *F*_(1, 45)_ = 18.16, *p* = 1.03 × 10^−4^; group: *F*_(1, 39)_ = 4.34, *p* = 0.044; Figure 6). This was true for isolated females when both layers and retainers w ere included in the analysis, and when only layers were considered (Figure S2).

**Figure 5.**
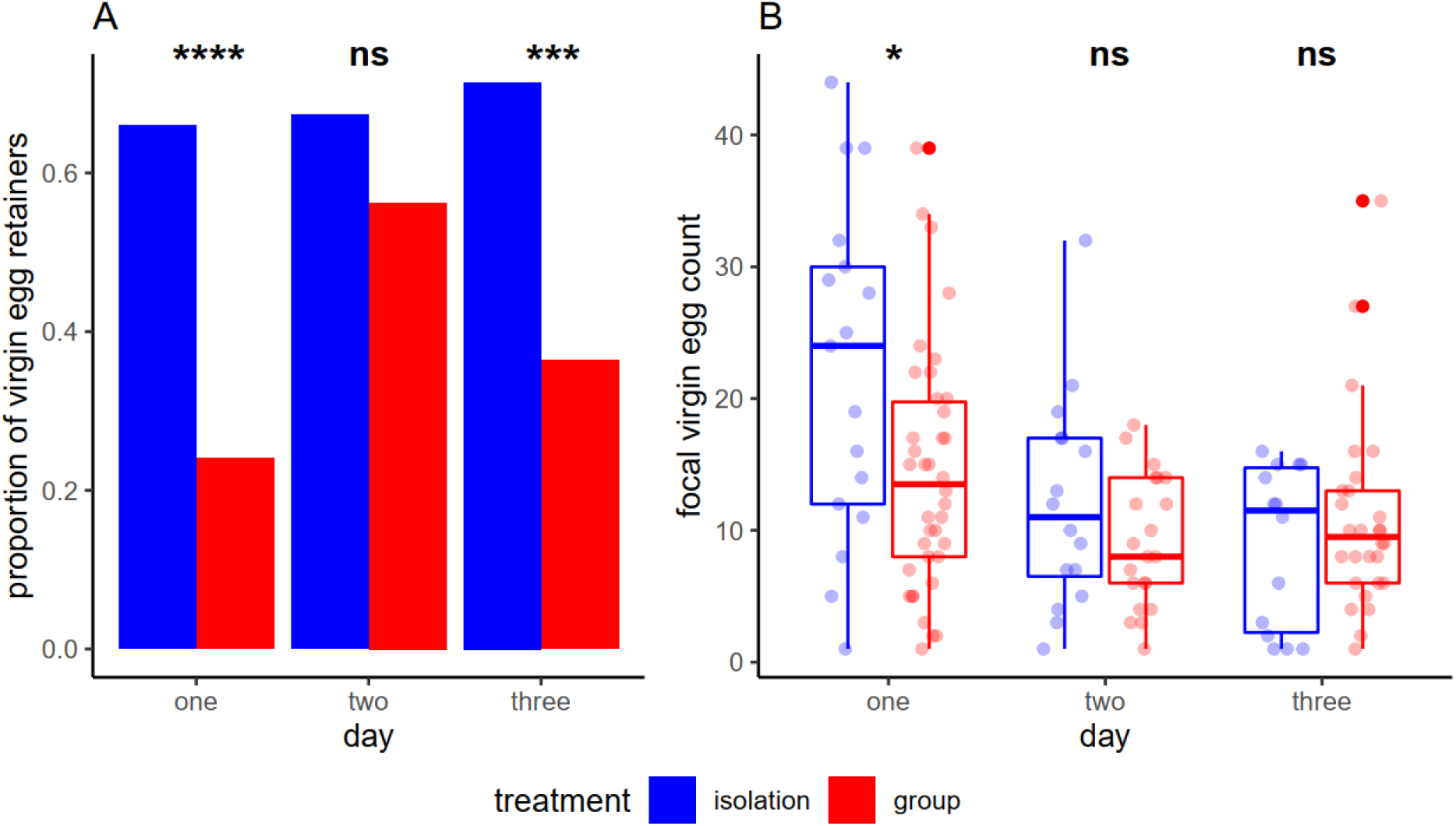
*D. melanogaster* females housed in isolation are more likely to retain virgin eggs. Virgin egg laying responses of *D. melanogaster* to the current social environment are shown. Focal females were kept in ‘isolation’ (blue bars/boxes) or ‘group’ (housed with three dyed non-focal females, red bars/boxes) treatments, for three days. (A) The proportion of female egg retainers (laying no eggs) on days one, two or three of social exposure. (B) Virgin egg counts of laying females (laying ≥ 1 egg on any given day) over three days of social exposure. Boxplots as in Figure 1.

**Figure 6.**
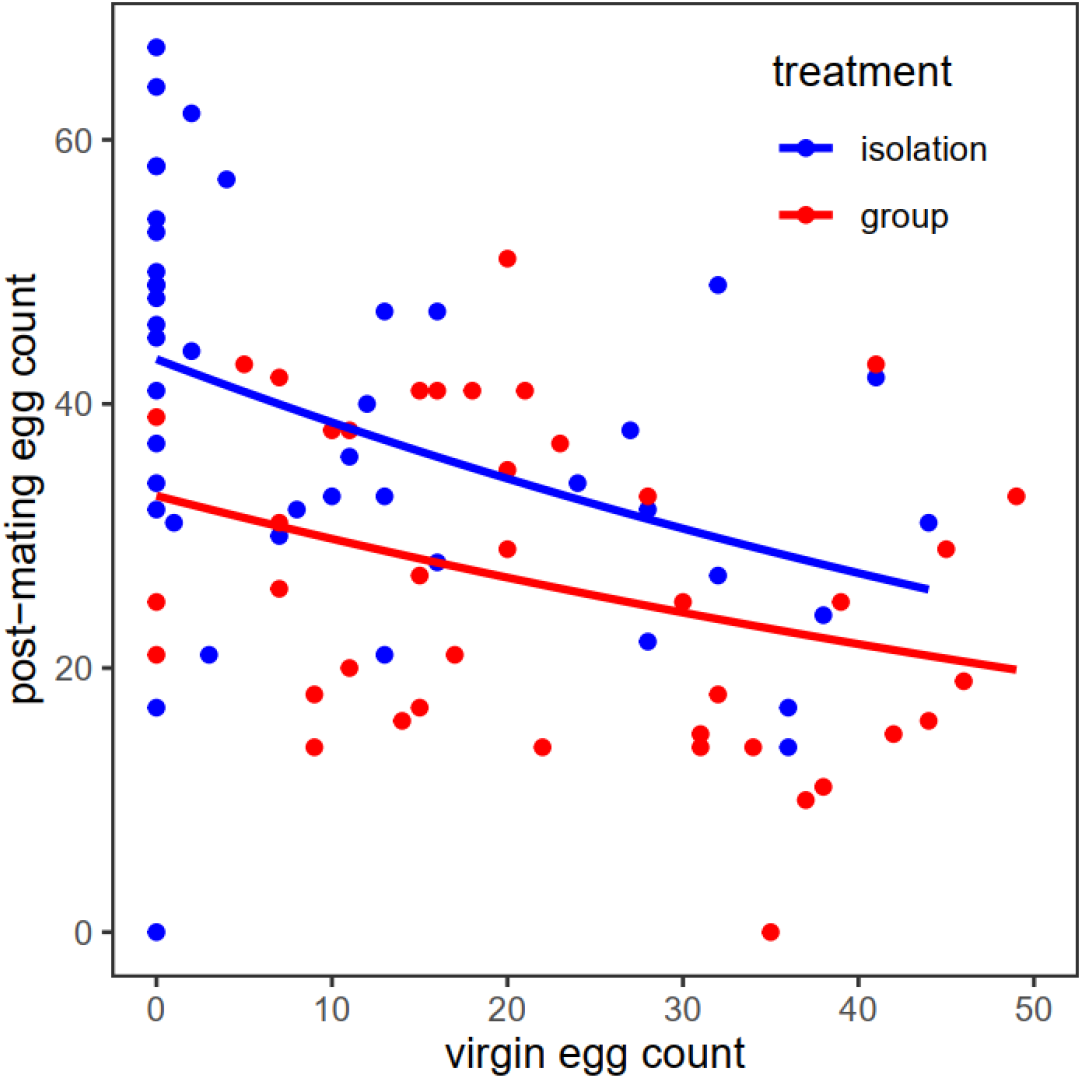
Negative relationship between pre- and post-mating fecundity in socially isolated and grouped females. Shown is the relationship between the total number of virgin eggs laid by a focal female in the three days prior to mating, and the number of post-mating eggs laid for 24h after mating. Focal females were held in either ‘isolation’ (blue) or in ‘group’ (with three Sudan red dyed non-focal females prior to mating, shown in red) treatments.

### Effect of social environment on mating latency and duration

Mating latency varied significantly with social environment in the control groups in five of the nine experiments (Figure S3, Table S3). In those five cases, previously grouped females were slower to mate than isolated females. Mating duration did not vary with social treatment in eight of the nine control experiments (Table S4). The exception was the 72h timepoint from the “length of social exposure” experiment in which previously grouped females had a significantly shorter mating duration than isolated females (Figure S4). Overall, there appeared to be no consistent effect of social exposure treatment on mating latency or mating duration.

## Discussion

The results show that female fecundity is strikingly plastic and varies according to the intrasexual social environment. Females exposed to groups of con - or heterospecific females in the pre-mating social environment showed significantly reduced post-mating fecundity compared to isolated females. Between 48-72h of exposure was required for fecundity to vary plastically. Direct contact with deposits left behind by previous females was sufficient to stimulate this plastic response, suggesting that the relevant cues are detected using tactile or gustatory pathways. Virgin egg retention was significantly higher among isolated in comparison to grouped females, leading to a negative relationship between virgin and post - mating fecundity, regardless of social treatment.

### Female fecundity varies plastically according to the con- and heterospecific social environment

The results reveal that the pre-mating social environment of female *D. melanogaster* significantly affects post-mating fecundity (see also Churchill *et al*., 2021). Such plasticity is expected to have profound fitness consequences for both the female experiencing the social environment and her mate. Females responding to others in their environment may gain benefits by optimising oviposition sites and food availability for offspring or through access to antimicrobials or anti-cannibalistic molecules deposited by other females or on the surface of eggs (Marchini *et al*., 1997; Narasimha *et al*., 2019). The presence of other adults and larvae at oviposition sites is known to have a significant impact on larval survival. Higher adult densities at oviposition sites lead to increased larval survival (Ashburner, 1989; Wertheim *et al*., 2002), likely through the suppression of fungal growth, but very high larval densities create competition and also lead to a lower larval survival rate (Wertheim *et al*., 2002). Therefore, a potential benefit of plasticity is that females adjust their oviposition rate in grouped situations to balance benefits of the suppression of microbial infection versus competition experienced by their larvae. The pattern we observed is consistent with potential benefits for grouped females in avoiding competition at oviposition sites by laying fewer eggs, and for isolated females to achieve density-dependent benefits by laying more. It is also possible that females alter their fecundity in order to benefit explicitly from the production of public goods. For example, in grouped situations, females might calibrate their fecundity to the level where they optimise benefits from the amount of tunnelling in the food medium and production of diffusible antimicrobials or anticannibalistic molecules (Marchini *et al*., 1997; Narasimha *et al*., 2019). Another explanation for grouped females laying fewer eggs after mating could be that they trade off offspring quantity for quality in environments where they expect their offspring to be in competition. It would be interesting to test for any such maternal effects by measuring offspring fitness traits.

Interestingly, the fecundity effect was not restricted to the conspecific social environment, as exposure of *D. melanogaster* females to either *D. simulans* or *D. yakuba* females also resulted in significantly reduced post-mating fecundity. Both *D. simulans* and *D. yakuba* are members of the *melanogaster* species subgroup, there is geographical overlap in the ranges of their populations, and all three species are generalists requiring rotting fruit for oviposition (Markow & O’Grady, 2005). The cues required for eliciting social responses may be conserved across this subgroup, with fecundity plasticity being triggered by the presence of any other females displaying these cues. Other types of sensory cues, such as chemical or pheromonal are known to be shared across closely related species. For example, aggregation pheromones across *D. melanogaster, yakuba* and *simulans* appear identical (Symonds & Wertheim, 2005) and attract heterospecifics as well as conspecifics in the field (Jaenike *et al*., 1992; Wertheim, 2001). There could be benefits to individuals from responding to cues emanating from heterospecifics if resources are shared and thus if the heterospecific cues signal resource quality or expected levels of competition for those limited resources. For example, larval resources may be exploited by several different species and so oviposition decisions based on the presence of heterospecifics could minimise over exploitation and have important fitness effects (Wertheim, 2005; Wertheim *et al*., 2002; Wertheim *et al*., 2002). We suggest that plasticity allows females to optimise their egg laying when oviposition and larval resources are likely to be utilised by closely-related species in sympatry. Interestingly, male *D. melanogaster* respond plastically to the presence of con- and some heterospecific males (*D. simulans* and *D. pseudoobscura*) but not others (*D. yakuba* or *D. virilis*) by increasing mating duration. However, the heterospecific responses when present do not occur to the same extent as following conspecific exposure (Bretman *et al*., 2017), likely because male responses to heterospecifics would carry costs but apparently little benefit (since heterospecifics pose minimal sperm competition). For females however, the consequences of basing oviposition decisions on the presence of heterospecifics or conspecifics may not differ markedly.

### Females require between 48-72h of social exposure to express fecundity plasticity

Responses by females to their social environments were not instantaneous, and appear to be longer than for the behavioural plasticity reported in males (Bretman *et al*., 2010). The precise social environment adult flies experience in the wild is likely to be subject to rapid changes, as flies eclose, move between patchy food resources or die. Such rapid variation may not provide a reliable indication of resource levels for females, thus setting up the requirement for a longer threshold of exposure to cues before decisions about potentially costly reproductive investment are triggered. Therefore, it is likely that the types of social responses seen in this study only benefit females if the social environment is sustained and thus accurately signals resource levels. We suggest that transient changes in social environment are unlikely to represent accurate indicators of resource quality to an even greater extent for females than males (Rouse & Bretman, 2016).

### Non-egg deposits from previous vial occupants stimulate the fecundity response

Interestingly, non-egg derived deposits left behind by other females were sufficient to stimulate post-mating fecundity responses. Of relevance is the observation that residual cues from either sex can also influence egg placement decisions in *D. melanogaster* (Malek & Long, 2020). Cues could include pheromones or microbes deposited from the cuticle or in the insect excreta (frass). Reproductively mature, virgin females harbour 50 types of cuticular hydrocarbon (CHC) and fatty acid molecules (Billeter & Wolfner, 2018). Female frass also contains CHCs such as methyl laurate, methyl myristate and methyl palmitate, and responses to deposited frass are reported to lead to increased feeding and aggregation (Keesey *et al*., 2016). Chemical cues are likely to be sensed by olfactory or gustatory sensory pathways, and indeed olfactory receptors were found to be partly responsible for behavioural changes in response to frass (Keesey *et al*., 2016). Frass deposits could provide a persistent and accurate indicator of the local population density and composition, and thus a more accurate indicator of potential resource levels as opposed to detection of the numbers of flies present at any given time, which could fluctuate rapidly.

### Direct contact with social cues is required, suggesting the use of gustatory sensory pathways

Females that were physically separated from other flies and eggs did not differ in fecundity from isolated females. Combined with our finding that non-egg derived female deposits are sufficient to stimulate plastic fecundity responses, these results suggest the gustatory (rather than tactile) pathways are used by females to respond to their social environment. Previous studies have found that female flies use sensory receptors located in their legs, ovipositor and proboscis to sample egg laying sites (Yang *et al*., 2008) and integrate olfactory and gustatory cues to make egg-laying decisions. Visual cues appeared not to be necessary; however, visually compromised *white* females did not exhibit fecundity plasticity. Possible explanations include pleiotropic effects of the *white* eye mutation such as impaired memory (Sitaraman *et al*., 2008), or compromised gravitaxis (Armstrong *et al*., 2006). That gustatory cues alone appear to be sufficient for females to assess and respond to social cues is in contrast to the multimodal strategy seen in males (Bretman *et al*., 2011). This may reflect the complexity of information required to make the appropriate response in each sex or the type of plastic phenotype involved.

### The social environment alters virgin egg retention

Isolated virgin females were more likely to retain eggs than those held in a group. This may be an adaptive strategy to conserve resources during long non-reproductive periods (Bouletreau-Merle & Fouillet, 2002) or when high quality oviposition sites are unavailable. Our finding that female *D. melanogaster* are more likely to retain virgin eggs in social isolation is consistent with observations for the tephritid *Rhagolettis pomanella* (Prokopy & Bush, 1973) and may indicate that a social stimulus is required for females to initiate ovulation. A benefit of high virgin egg retention was increased fecundity following mating, consistent with previous findings (Edward *et al*., 2014).

### Mating behaviour was not consistently affected by social environment in females

The effects of social exposure on mating latency were inconsistent, as is also found in males (Bretman *et al*., 2009; Bretman *et al*., 2013; Bretman *et al*., 2013; Dore *et al*., 2020). Individuals may be differentially susceptible to environmental differences between experiments or changing population dynamics in the stock cages from which they were collected. In almost all cases mating duration was unaffected by female social environment. This contrasts with the corresponding plasticity seen in males (Bretman *et al*., 2009) and reflects the finding that mating duration is largely under male control (Bretman *et al*., 2013). Additionally, it suggests that males do not respond to the social environment of their mate despite potential fitness costs if the female has lowered fecundity.

## Conclusions

These results represent a significant advance in knowledge of how the intrasexual social environment affects female reproduction. We investigated responses to both con- and heterospecifics, the length of exposure required to express plasticity, and the cues and mechanisms underlying the fecundity response. We found that the social environment does indeed have the potential to affect female fitness. A key, important outcome is that the responses, timing and nature of cues used are markedly different in females vs males, and this likely reflects the contrasting benefits of reproductive plastic behaviour between the sexes.

## Methods

### Fly stocks and handling

Wild type *D. melanogaster* flies were from a large laboratory population originally collected in the 1970s in Dahomey (Benin) and maintained in stock cages with overlapping generations. Wild type *D. simulans* and *D. yakuba* were obtained from the San Diego *Drosophila* Stock Center and KYORIN-Fly *Drosophila* species stock centre (stock #k-s03), respectively. Flies were reared on standard sugar yeast (SY) medium (100 g brewer’s yeast, 50 g sugar, 15 g agar, 30 ml Nipagin (10% w/v solution), and 3 ml propionic acid, per litre of medium) in a controlled environment (25°C, 50% humidity, 12:12 hour light:dark cycle). For the Sudan Red food medium, 800 ppm Sudan Red 7B (*Sigma Aldrich*) dye was added to the SY diet before dispensing. Eggs were collected from population cages on grape juice agar plates (50 g agar, 600 ml red grape juice, 42 ml 10% w/v Nipagin solution per 1.1 l H_2_O) supplemented with fresh yeast paste, and first instar larvae were transferred to SY medium at a standard density of 100 per vial (glass, 75×25mm, each containing 7ml medium). Male and female adults were separated within 6h of eclosion under ice anaesthesia and stored in single sex groups of 10/vial. *White* females were from a stock carrying the *w*^*1118*^ allele that had been backcrossed three times into the Dahomey wild type. *Orco* females were generated from backcrossing *Orco*^*1*^ (Bloomington Drosophila Stock Centre, stock #23129) stock for three generations into a Dahomey stock carrying the *TM3 sb ry* balancer on chromosome 3. Eggless females were generated by crossing males from the *Ovo*^*D1*^ stock (Bath *et al*., 2017) with wild type Dahomey females.

### Effect on female mating behaviour and fecundity of variation in pre-mating social environment

In all experiments, virgin focal *D. melanogaster* females were CO_2_ anaesthetised at 3-4 days old and assigned to isolation (1 female per vial) or group (1 focal and 3 virgin non-focal females per vial) social treatments. Females were exposed to these social environments for a period of 72h (unless stated otherwise) prior to mating. Wildtype males were aspirated individually into fresh SY vials the day prior to the mating trial. Mating trials were conducted at 25°C at 50% RH, always starting at 9 am in the morning unless otherwise stated. On the day of mating, focal females were aspirated into vials containing a single male. Pairs were observed and the introduction time, start and end of mating were recorded. Any flies that did not start mating within 90 min were discarded. Males were removed immediately following the end of copulation and females left to oviposit for 24h before being discarded. Eggs laid on the surface of the SY medium in this 24h period were counted under a Leica MZ7.5 stereomicroscope. Sample sizes for all experiments are shown in Table S1.

### Female fecundity responses to variation in the social environment and effect of exposure to con- vs hetero-specific females

Following the protocol as described above, focal wildtype *D. melanogaster* females were kept in isolation or housed with 3 non-focal females of the same or two different *Drosophila* species. We chose as heterospecific treatments two species of the *melanogaster* subgroup - *D. simulans* and *D. yakuba*, which shared their last common ancestor with *D. melanogaster* ∼5 MYA and ∼13 MYA, respectively (Tamura *et al*., 2004). Non-focal females were wing - clipped under CO_2_ anaesthesia prior to setting up the social exposure treatments, in order to distinguish them from the focal *D. melanogaster* individuals.

### Effect of length of social exposure period on post-mating fecundity

The experiment was set up following the standard protocol above, with wildtype Dahomey focal and non-focal females, but with varying lengths of social exposure before mating. To test the effect on post-mating female fecundity from shorter term exposure, all females were placed into the social environments in parallel (between 9 and 10am on the day of the mating trails), then subsets of focal females were mated after 2, 4 or 8h. Therefore, these matings were conducted at different times of the day (2h at 12pm, 4h at 2pm, and 8h at 6pm). Longer-term exposure was tested in a separate experiment. Again, all social environments were set up in parallel, then mating trials on subsets of focal females were conducted after 24, 48 and 72h, all at 9am each day.

### Investigation of whether exposure to eggs or to female deposits in the absence of eggs are required for social exposure effects on post-mating fecundity

This experiment was carried out in two sets. In the first, we tested whether exposure to eggs of other females, or deposits of other females in the absence of eggs, were required for females to show plastic fecundity responses after mating. To do this we used non-focal females from the *Ovo*^*D1*^ (eggless) genotype. Wildtype focal females were kept alone (isolation), exposed to 3 wildtype non focal conspecifics (group), 3 eggless *Ovo*^*D1*^ non-focal females (group - eggless females), or to an SY vial that had previously housed 3 eggless *Ovo*^*D1*^ females for the preceding 24h (isolation - female deposits). In the second set, wildtype focal females were again kept alone (isolation), exposed to 3 wildtype non focal conspecifics (group) or exposed to eggs laid in the previous 24h by three wildtype non-focals (isolation - egg-spiked). In both experiment sets, all focal females were moved to “fresh” (deposits, egg-spiked or clean food) vials every 24h of the exposure period to maintain the strength of the specific cues involved.

### Investigation of the sensory pathways required to detect cues of social exposure effects on post-mating fecundity

To identify the sensory pathways used by females to detect the proxies of female presence described above, we conducted three sets of experiments, each with standard isolation and group control treatments. To test the effect on post mating fecundity of manipulating visual inputs, we used either wildtype females held in darkness, or visually-defective *white* focal females held under normal light conditions (Ferreiro *et al*., 2018). Non-focal females were all wildtype. To test the effect of manipulating olfactory cues we used focal females with a knockout mutation in the *Orco* gene (encoding a broadly expressed odorant receptor, essential for olfaction of a wide range of stimulants (Larsson *et al*., 2004)), or we surgically removed the third antennal segment of wildtype focal females under CO_2_ anaesthesia one day prior to setting up the social treatments. The antennal segment contains sensillae bearing odorant receptors, but also aristae that detect sound (Göpfert & Robert, 2001; van der Goes van Naters & Carlson, 2007). Non-focal females for both olfactory experiments were wildtype females with intact antennae, which were wing-clipped under CO2 anaesthesia one day prior to social exposure. Finally, to test the effect of manipulating tactile cues, we physically separated wildtype focal females from non-focals using a perforated acetate divider to create two chambers within a standard vial. Perforations allowed the transmission of sound and odours, and the dividers were translucent which allowed for the perception of visual cues.

### Effect of social environment on virgin egg retention

In the final experiment we used a novel egg marking procedure to test the effect of isolation and group treatments on pre-mating (virgin) egg production and retention. Wild type focal females were reared according to the standard protocol. Non-focal females were reared from the 1^st^ instar larval stage on SY food containing 800 ppm oil-based Sudan Red dye, which stains lipids, resulting in the production and laying of visibly pink eggs as adults. Dyed females were collected upon eclosion and maintained on Sudan Red food for 3-4 days prior to setting up the social treatments. Social treatments were set up according to the standard protocol, above. For the group treatment, one focal female was housed in a vial with three dyed non-focals. Females were then moved every 24h to fresh food until mating. The number of white and dyed (pink) eggs laid by the focal and non-focal females, respectively, was recorded for each 24h period of social exposure. Mating trials and post-mating egg counts were conducted as above.

### Statistical analysis

Statistical analyses were carried out in R v 3.6.3 (R Core Team, 2013). Post-mating egg counts were analysed using a generalised linear model (GLM) with a log link and quasi - Poisson errors to account for over-dispersion. The total number of virgin ‘egg layers’ (females that laid ≥ 1 egg on a given day) versus ‘retainers’ (no eggs laid on a given day) in each social treatment was analysed using a Chi-square test. The number of virgin eggs laid by ‘egg-layers’ (non-zero counts) across social treatments was analysed using a GLM with quasi-Poisson errors. Significance values for GLMs were derived from an anova F test of the model. Mating latency was analysed using Cox Proportional Hazards models, fitted using the “coxph” function from the “survival” package. Individuals that did not mate within 90 minutes were treated as censors. For mating duration, times of < 6 min and > 30 min were excluded from the analysis. These data points represent extremely short copulations, in which genitalia were unlikely to have been fully engaged or sperm transferred (Gilchrist & Partridge, 2000). Very long copulations can result if genitalia become “stuck” and flies fail to disengage. In total, 11 such outliers were removed from across five of the mating duration experiments (supplementary table S2). Mating duration data were normally distributed for each experiment (Shapiro-Wilk tests, p > 0.05) and were analysed using Welch two sample *t*-tests.

## Authors’ contributions

EKF, AB and TC conceived the study, EKF, SL, WR and AT conducted the experiments and analyses, EKF analysed the data and EKF, SL and TC wrote the paper. All authors read and approved the final version of the manuscript.

## Acknowledgements

We thank Jean-Christophe Billeter for helpful comments on the manuscript, Alice Dore, Nick West, Nathan McConnell, Lucy Friend, Mike Darrington and Jessy Rouhana for help with the mating assays, Paul Candon and Kerri Armstrong for technical assistance and Ellie Bath for sending us the *OvoD1* strain.

## Statement on data sharing

All raw data will be made available on the DRYAD data repository upon acceptance. We will also provide a private data sharing link to the raw data, if requested by the reviewers.

