## Supplementary material for "Female fruitflies use gustatory cues to exhibit reproductive plasticity in response to the social environment"

**Table S1.** Treatment sample sizes for post-mating eggs, mating latency and mating duration response variables across all experiments. Mating latency was analysed using a cox proportional hazards test with censorship and the number of censored points included in the overall sample size are indicated in brackets with an asterisk. For mating duration, the final sample size is given following the removal of outlying points if applicable (durations of < 6 minutes and > 30 minutes). The numbers of outlying points removed in each case are indicated in brackets.

| Experiment | Treatment | post-mating eggs | mating latency (*censored points) | mating duration (number of removed outliers) |
|---|---|---|---|---|
| 1 (baseline) | isolation | 46 | 47 (1*) | 46 (1) |
| | group | 40 | 40 (4*) | 37 (3) |
| 2 (temporal) | **isolation** | | | |
| | 2 hrs | 43 | - | - |
| | 4 hrs | 44 | - | - |
| | 8 hrs | 35 | - | - |
| | 24 hrs | 58 | - | - |
| | 48 hrs | 57 | - | - |
| | 72 hrs | 60 | 61 | 61 |
| | **group** | | | |
| | 2 hrs | 46 | - | - |
| | 4 hrs | 44 | - | - |
| | 8 hrs | 42 | - | - |
| | 24 hrs | 59 | - | - |
| | 48 hrs | 60 | - | - |
| | 72 hrs | 62 | 62 (1*) | 62 |
| 3 (heterospecific) | isolation | 46 | - | - |
| | *simulans* | 32 | - | - |
| | *yakuba* | 46 | - | - |
| 4a (eggless) | isolation | 44 | 44 (6*) | 44 |
| | group | 39 | 40 (7*) | 40 |
| | eggless | 35 | - | - |
| | conditioned | 46 | - | - |
| 4b (spiked egg) | isolation | 39 | 39 | 39 |
| | group | 39 | 39 | 39 |
| | egg-spiked | 32 | - | - |
| 5a (olfactory, antennaless) | **control** | | | |
| | isolation | 43 | 44 (2*) | 44 |
| | group | 42 | 42 (4*) | 41 (1) |
| | **antennaless** | | | |

| | | | | |
|---|---|---|---|---|
| | isolation<br>group | 35<br>29 | -<br>- | -<br>- |
| 5b (olfactory, | **control** | | | |
| *Orco*) | isolation | 48 | 48 (2*) | 48 |
| | group | 42 | 42 (5*) | 41 (1) |
| | ***Orco*** | | | |
| | isolation | 32 | - | - |
| | group | 36 | - | - |
| 5c<br>(tactile/gustatory) | **control**<br>isolation<br>group<br>**divided**<br>isolation<br>group | <br>41<br>42<br><br>48<br>38 | <br>42 (2*)<br>42 (2*)<br><br>-<br>- | <br>39 (3)<br>42<br><br>-<br>- |
| 5d (visual) | **control**<br>isolation<br>group<br>**dark**<br>isolation<br>group<br>**white**<br>isolation<br>group | <br>43<br>41<br><br>43<br>45<br><br>48<br>41 | <br>43 (2*)<br>42 (2*)<br><br>-<br>-<br><br>-<br>- | <br>43<br>42<br><br>-<br>-<br><br>-<br>- |
| 6 (sudan red) | isolation<br>group | 47<br>41 | 46 (2*)<br>41 (1*) | 46<br>39 (2) |

**Table S2.** Chi-squared analysis of the number of egg layers (laying ≥ 1 virgin egg) versus number of egg retainers (zero eggs laid) for virgin females held in isolation or housed in groups of four for 3 days. Number of virgin eggs laid by the focal females were counted each day for three days. Data were also analysed for the three days combined.

| | **Day 1** | | | **Day 2** | | | **Day 3** | | |
|---|---|---|---|---|---|---|---|---|---|
| treatment | layers | retainers | Chi square | layers | retainers | Chi square | layers | retainers | Chi square |
| isolation | 17 | 33 | $X^2 = 17.8$ *df* = 1 $p =$ 2.43e-05 | 16 | 33 | $X^2 =$ 1.3 *df* = 1 $p =$ 0.26 | 14 | 35 | $X^2 =$ 11.5 *df* = 1 $p =$ 0.0007 |
| group | 38 | 12 | | 21 | 27 | | 28 | 16 | |

| | **All days combined** | | |
|---|---|---|---|
| treatment | layers | retainers | Chi square |
| isolation | 29 | 20 | $X^2 = 12.2$ *df* = 1 $p =$ 0.00048 |
| group | 40 | 4 | |

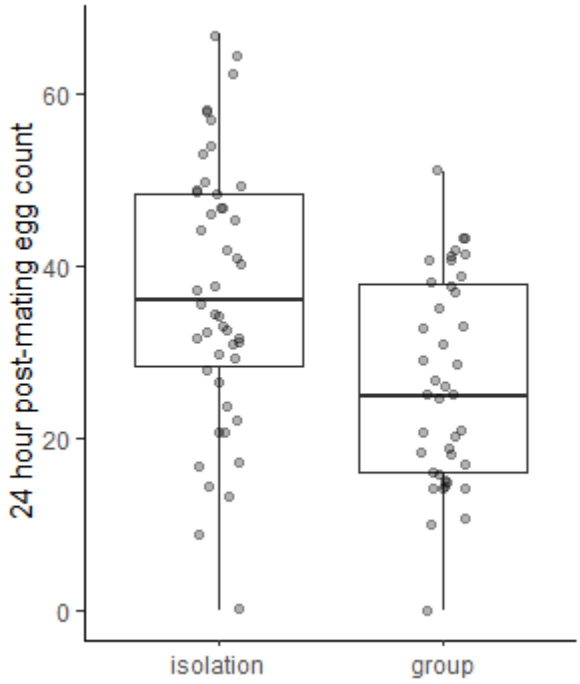


**Figure S1.** Effect of pre-mating social environment on post-mating fecundity for focal females exposed to Sudan red dyed non-focal females in the group treatment.

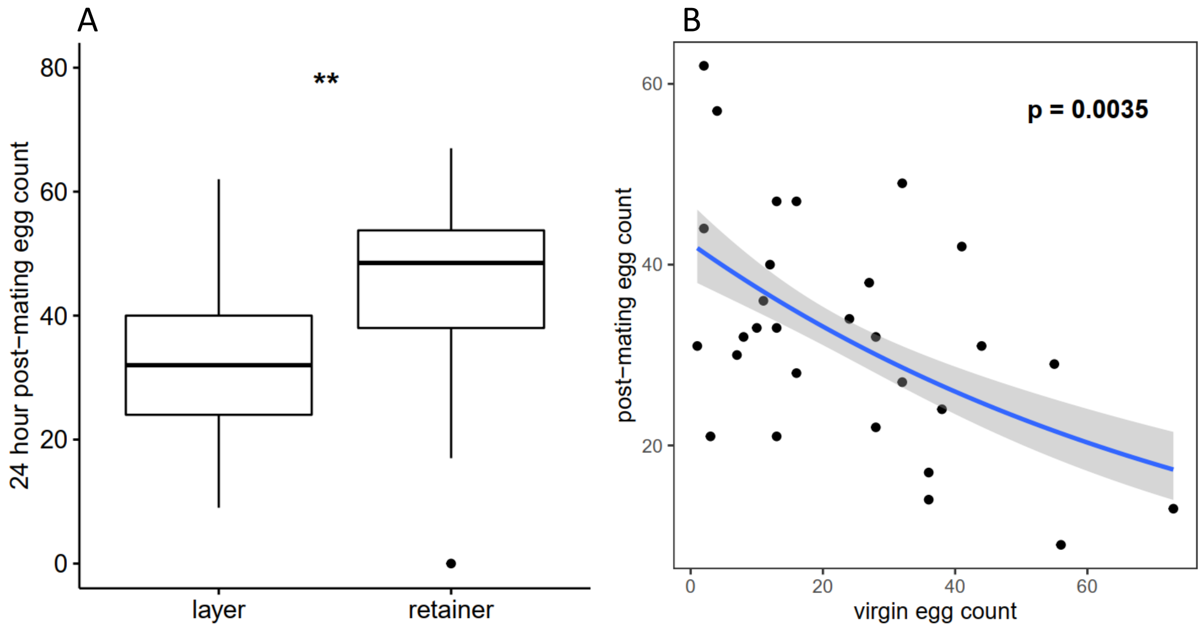


**Figure S2.** Effect of number of virgin eggs laid over three days on post-mating fecundity by females held in isolation prior to mating. (A) Post-mating egg count was compared for females defined as layers (≥ 1 virgin egg laid) and retainers (zero virgin eggs laid) ($F_{(1,45)} = 7.91$, $p = 0.007$). (B) Relationship between virgin and post-mating eggs for layers only ($F_{(1,27)} = 10.94$, $p = 0.003$).

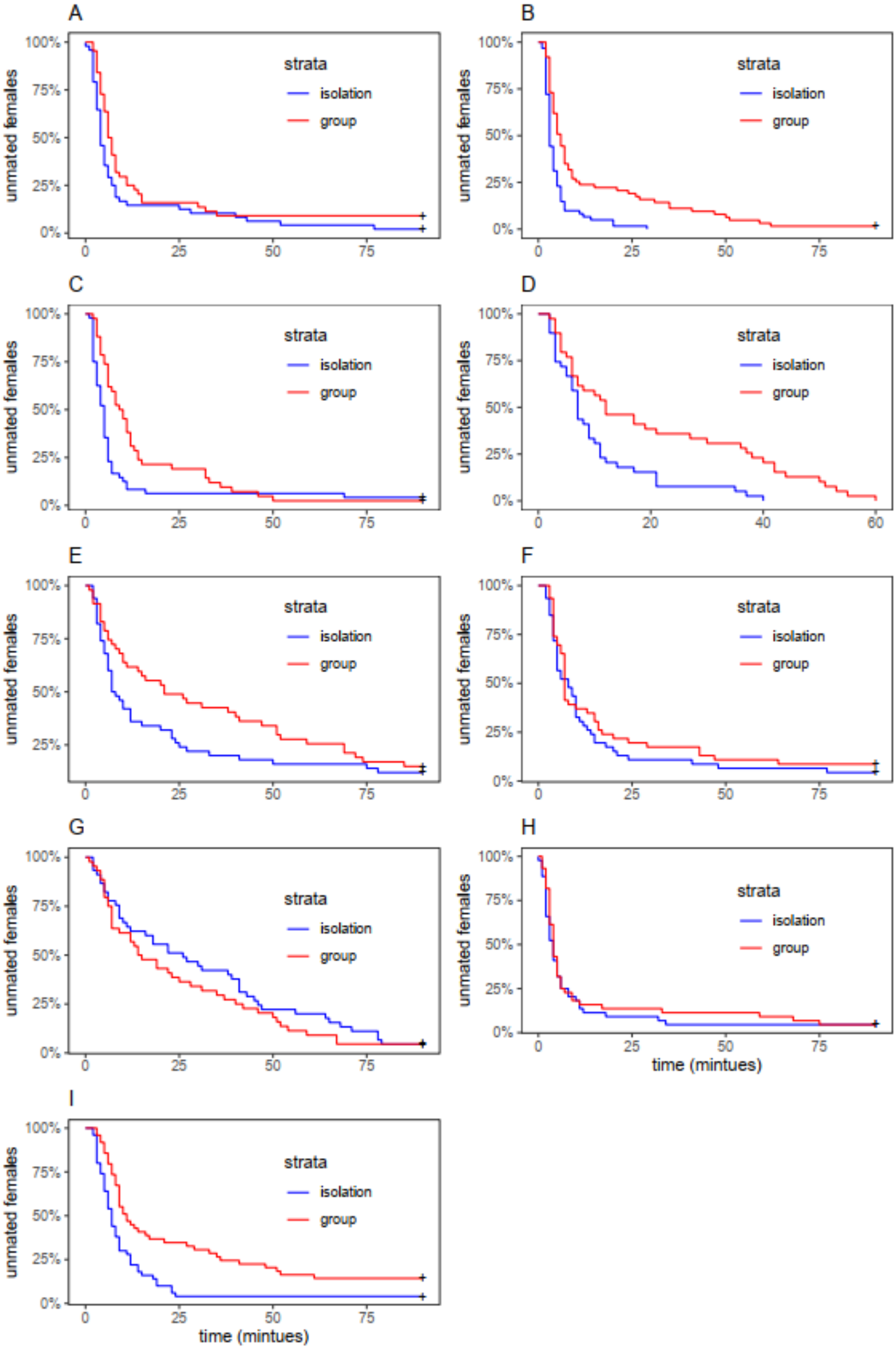
A
B
C
D
E
F
G
H
I
strata
isolation
group
unmated females
time (mintues)
100%
75%
50%
25%
0%
0
25
50
75
20
40
60

**Figure S3.** Effect of social environment on mating latency across eight separate experiments in this study. Females were either kept in isolation (blue) or housed in groups of four (red) for 72 hours prior to mating. The experiments were: (A) baseline responses (Experiment 1 in the main text); (B) Effect of length of exposure to pre-mating social environment, 72 hr timepoint only (Experiment 2 in main text); (C) Effect of social environment on virgin eggs (Experiment 6 in main text); (D) control from the "egg-spiked" block (Experiment 4 in main text); (E) control from the "OvoD1" block (Experiment 4 in main text); (F) control from the olfactory cues experiment; (G) control from the visual cues experiment; (H) control from the tactile cues experiment; (I) control from the *Orco1* experiment (all Experiment 5 in main text).

**Table S3.** Cox proportional hazards analysis output for effect of social environment on mating latency.

| Experiment | z | p-value | lower 95% CI | upper 95% CI |
|---|---|---|---|---|
| A (baseline) | -2.03 | 0.04 | 0.42 | 0.99 |
| B (exposure length) | -4.36 | 1.32e-05 | 0.29 | 0.63 |
| C (Sudan red) | -2.96 | 0.0031 | 0.34 | 0.80 |
| D (egg spiked) | -3.20 | 0.0014 | 0.27 | 0.73 |
| E (eggless) | -1.64 | 0.10 | 0.45 | 1.07 |
| F (olfactory) | -0.89 | 0.38 | 0.54 | 1.26 |
| G (visual) | 1.05 | 0.29 | 0.83 | 1.93 |
| H (tactile/gustatory) | -0.60 | 0.55 | 0.57 | 1.35 |
| I (*orco1*) | -3.38 | 0.00072 | 0.31 | 0.73 |

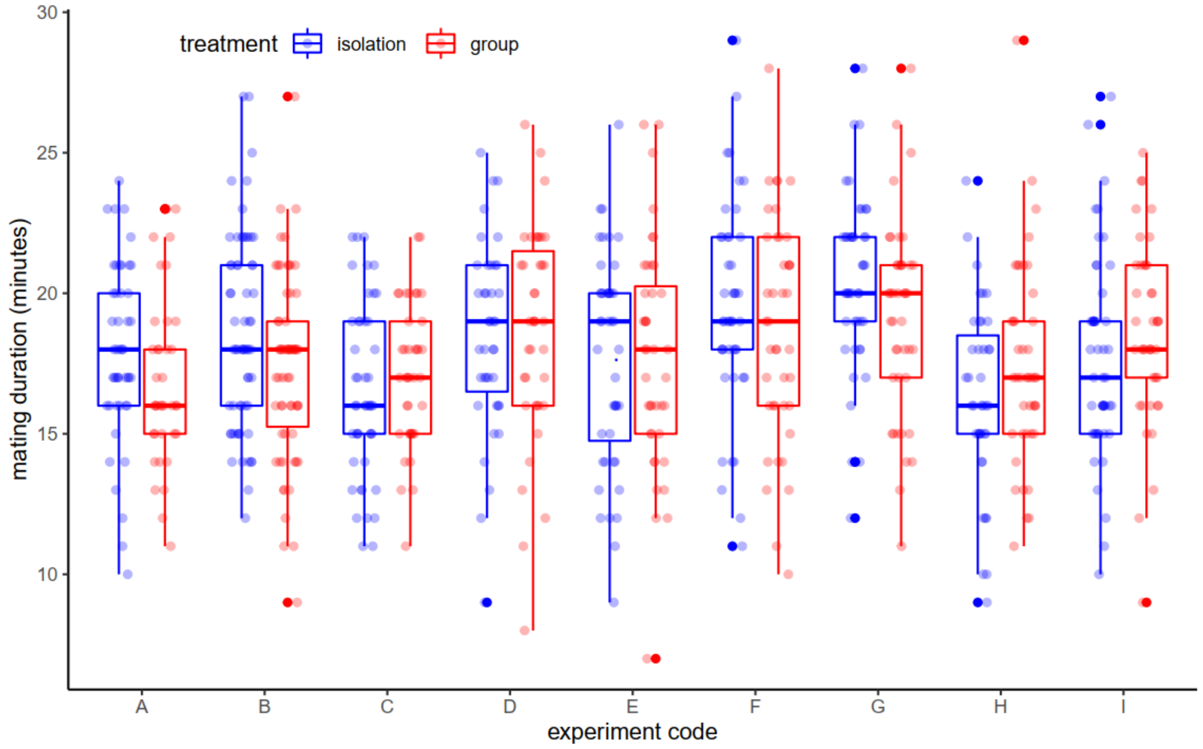


**Figure S4.** Mating duration of females kept in social-isolation or in a group prior to mating across eight separate experiments in this study. For each experiment, only the data for control flies was analysed to enable direct comparison between experiments (i.e. all data are for intact, wildtype *D. melanogaster* females set up according to the standard experiment protocol). The experiments were: (A) baseline responses (Experiment 1 in the main text); (B) Effect of length of exposure to pre-mating social environment, 72 hr timepoint only (Experiment 2 in main text); (C) Effect of social environment on virgin eggs (Experiment 6 in main text); (D) control from the “egg-spiked” block (Experiment 4 in main text); (E) control from the “OvoD1” block (Experiment 4 in main text); (F) control from the olfactory cues experiment; (G) control from the visual cues experiment; (H) control from the tactile cues experiment; (I) control from the *Orco1* experiment (all Experiment 5 in main text).

**Table S4.** Effect of social environment on mating duration. Output from Welch's two-sample t-test.

| Experiment | mean duration (minutes) | t | df | p-value | lower 95% CI | upper 95% CI |
|---|---|---|---|---|---|---|
| A (baseline) | I = 17.9<br>G = 16.8 | 1.62 | 78.9 | 0.11 | -0.25 | 2.44 |
| B (exposure length) | I = 18.6<br>G = 17.3 | 2.08 | 120.5 | 0.04 | 0.06 | 2.48 |
| C (Sudan red) | I = 16.5<br>G = 17.1 | -1.03 | 82.9 | 0.31 | -1.84 | 0.58 |
| D (egg spiked) | I = 18.5<br>G = 18.6 | -0.22 | 74.8 | 0.83 | -1.81 | 1.45 |
| E (eggless) | I = 17.7<br>G = 17.8 | -0.08 | 80.2 | 0.94 | -1.77 | 1.63 |
| F (olfactory) | I = 19.5<br>G = 18.7 | 0.91 | 81.7 | 0.36 | -0.91 | 2.45 |
| G (visual) | I = 20.4<br>G = 19.1 | 1.78 | 81.8 | 0.08 | -0.15 | 2.75 |
| H (tactile/gustatory) | I = 16.4<br>G = 17.4 | -1.40 | 79.0 | 0.16 | -2.59 | 0.45 |
| I (*orco1*) | I = 17.7<br>G = 18.6 | -1.26 | 86.5 | 0.21 | -2.37 | 0.53 |
